# Novel CRISPR/Cas9 gene drive constructs in *Drosophila* reveal insights into mechanisms of resistance allele formation and drive efficiency in genetically diverse populations

**DOI:** 10.1101/112011

**Authors:** Jackson Champer, Riona Reeves, Suh Yeon Oh, Chen Liu, Jingxian Liu, Andrew G. Clark, Philipp W. Messer

**Affiliations:** Department of Biological Statistics and Computational Biology; Department of Molecular Biology and Genetics, Cornell University, Ithaca, NY 14853

**Author notes:** Corresponding authors: JC, PWM.

## Abstract

A functioning gene drive system could fundamentally change our strategies for the control of vector-borne diseases by facilitating rapid dissemination of transgenes that prevent pathogen transmission or reduce vector capacity. CRISPR/Cas9 gene drive promises such a mechanism, which works by converting cells that are heterozygous for the drive construct into homozygotes, thereby enabling super-Mendelian inheritance. Though CRISPR gene drive activity has already been demonstrated, a key obstacle for current systems is their propensity to generate resistance alleles. In this study, we developed two CRISPR gene drive constructs based on the *nanos* and *vasa* promoters that allowed us to illuminate the different mechanisms by which resistance alleles are formed in the model organism *Drosophila melanogaster.* We observed resistance allele formation at high rates both prior to fertilization in the germline and post-fertilization in the embryo due to maternally deposited Cas9. Assessment of drive activity in genetically diverse backgrounds further revealed substantial differences in conversion efficiency and resistance rates. Our results demonstrate that the evolution of resistance will likely impose a severe limitation to the effectiveness of current CRISPR gene drive approaches, especially when applied to diverse natural populations.

## INTRODUCTION

Gene drive systems promise a mechanism for rapidly spreading alleles in a population through super-Mendelian inheritance^1–5^. One prominent example is the homing drive, in which the drive allele contains an endonuclease that targets a specific site in the genome for cleavage and then inserts itself into that site via homology-directed repair (HDR). A heterozygote for the drive allele can thereby be converted into a homozygote, resulting in most its progeny inheriting the drive allele. In principle, this allows for the rapid spread of such an allele in the population even if it carries a fitness cost to the organism^6–9^.

Via this mechanism, a genetic payload could be rapidly disseminated throughout an entire population^6–11^, opening up a variety of fascinating potential applications. For example, a functioning gene drive could provide a highly efficient means for controlling vector borne diseases such as malaria^1–5^, which kills over 400,000 people per year^12^ and is notorious for rapidly acquiring drug resistance to every newly-deployed drug, including the current frontline drug Artemisinin^13,14^. Genetic payloads for reducing malaria transmission in mosquitoes have already been successfully tested^15–17^. Combined with a gene drive, they could provide a promising new approach to fight this devastating disease. Other proposed applications range from spreading genetically engineered antiviral effector genes against dengue^18^, to suppressing the populations of invasive crop pests such as *Drosophila suzukii*^19^.

Early attempts at constructing a homing gene drive were based on *I-SceI* and *I-OnuI* enzymes with artificial target sites in the fruit fly *Drosophila melanogaster^20–23^* and the mosquito *Anopheles gambiae^24^.* These approaches had rather limited success, primarily because of low drive conversion rates. More recently, researchers have utilized the CRISPR/Cas9 system for homing drives with much higher efficiency in *D. melanogaster^25^* and *Saccharomyces cerevisiae* yeast^26^. The design of such CRISPR gene drives has now also been demonstrated in mosquitoes, including approaches aimed for population suppression of *A. gambiae^27^* and for spreading a malaria resistance gene in *Anopheles stephensi^28^*. These approaches use a Cas9 endonuclease to cleave chromosomes at a specific location dictated by a guide RNA (gRNA), which can be engineered to target a wide range of nucleotide sequences.

One of the key obstacles to a successful gene drive approach lies in the emergence of resistance against the drive^29,30^. Such resistance is particularly pertinent to CRISPR gene drives, as they are expected to produce resistance alleles themselves when Cas9-induced cleavage is repaired by non-homologous end-joining (NHEJ), instead of drive incorporation by HDR. NHEJ will often result in mutated target sites that are no longer recognized by the drive’s gRNA ^25,27,28^.Theoretical studies have shown that the formation of such resistance alleles severely limits the ability of a drive to spread in large populations^29,30^.

All CRISPR gene drive constructs tested to date in insects have produced resistance alleles in significant quantities. This phenomenon was probably best assessed in a study in *A. stephensi*^28^, which found a very high rate (~98%) of successful conversion of wild type alleles into drive alleles in the germline of heterozygotes, but also a high rate (>77%) at which resistance alleles were formed post-fertilization in embryos produced by females with the drive, presumably due to persistence of maternal Cas9 or "leaky" expression^28^. Such high rates of resistance allele formation would almost certainly prevent the spread of the drive allele in any wild population, especially if it carries a fitness cost^29,30^.

It is clear that to establish CRISPR gene drive as a practical means for genetic transformation of natural populations, the propensity to generate resistance alleles needs to be substantially reduced. Achieving this goal requires a better understanding of the mechanisms by which resistance alleles arise and the factors that determine these processes. Importantly, we must include the possibility that drive efficiency and rates of resistance allele formation might vary between individuals in the population, for instance when genetic variability affects expression levels of Cas9, or the efficiency and fidelity of different cleavage-repair pathways. Even if the formation of resistance alleles could be effectively suppressed in most individuals of a population, some individuals with high rates of resistance allele formation would likely thwart a drive in the long-term.

In this study, we use two newly-developed gene drive constructs in the model organism *D. melanogaster* to quantify drive efficiency as well as rates and mechanisms of resistance allele formation. One of our constructs resembles the *vasa* promoter-driven construct originally developed by Gantz and Bier^25^, with the addition of a dsRed fluorescent protein as payload that allows us to easily detect driver alleles in heterozygotes. The other uses the *nanos* promoter, which is known to have germline-restricted expression with lower toxicity than *vasa*^31^ and may represent a better alternative to express Cas9. We further study drive parameters and resistance allele formation rates of these constructs in genetically distinct backgrounds, including flies from the Global Diversity Lines from five continents^32^.

## MATERIALS & METHODS

### Plasmid construction

Plasmids were constructed using standard molecular biology techniques based on Gibson Assembly Master Mix (New England Biolabs) and JM109 competent cells (Zymo Research). Restriction enzymes were from New England Biolabs (except *Fsp*AI, from Thermo Scientific). Miniprep, gel extraction, and other DNA purification kits were from Zymo Research. PCR was conducted with Q5 Hot Start DNA Polymerase (New England Biolabs) according to the manufacturer’s protocol. The Supplementary Methods contain a list of plasmids generated in this study and a list of DNA oligo sequences used to construct and sequence these plasmids and genomic gRNA target sites. Plasmids pDsRed-attP (Addgene plasmid #51019) and pvasa-Cas9^33^ were provided by Melissa Harrison, Kate O'Connor-Giles, and Jill Wildonger. Plasmids pCFD3-dU6:3gRNA^31^ (Addgene plasmid #49410) and pnos-Cas9-nos^31^ (Addgene plasmid #62208) were provided by Simon Bullock. Cas9 gRNA target sequences were identified by the use of CRISPR Optimal Target Finder^33^.

### Generation of transgenic lines

To transform a *w^1118^ D. melanogaster* line with a homing drive, the primary donor plasmid (either IHDyN1 or IHDypV1) was purified using a ZymoPure Midiprep kit (Zymo Research). A supplemental source of Cas9 from plasmid pHsp70-Cas9^34^ (provided by Melissa Harrison & Kate O'Connor-Giles & Jill Wildonger, Addgene plasmid #45945), was included in the injection to ensure cleavage of the target site, as was a supplemental source of the gRNA (IHDyg1 or IHDypg1) targeting the insertion site at the X-linked *yellow* gene. Concentrations of donor, Cas9, and gRNA plasmids were approximately 138, 45, and 18 ng/μL, respectively, in 10 mM tris-HCl, 23 μM EDTA, pH 8.1 solution. Injections were performed by GenetiVision into a *w^1118^* line. Insertion of the donor plasmid was confirmed by rearing injected G0 embryos to adulthood and crossing them with *w^1118^* flies. G1 progeny were then screened for the presence of dsRed fluorescent protein in their eyes, which was indicative of successful transformation. dsRed fluorescent flies were then crossed together until all male progeny were dsRed fluorescent for two consecutive generations, indicating that the stock was homozygous for the drive allele.

### Fly rearing and phenotyping

Flies were reared at 25°C on Bloomington Standard media with a 14/10 hour day/night cycle. For general maintenance, stocks were provided with new food every 2-3 weeks. Flies were anesthetized by CO2 during phenotyping. Yellow phenotype was assessed in the body and wings, and flies were considered “mosaic” if they displayed any visible level of mosaicism on any part of their body or wings. To assess fluorescent red phenotype conferred by the dsRed transgene, the NIGHTSEA system (NIGHTSEA) was used with a conventional stereo dissecting microscope. All work with live gene drive flies was performed at the Sarkaria Arthropod Research Laboratory at Cornell University, a USDA APHIS-inspected Arthropod Containment Level 3 (ACL-3) insect quarantine facility. Strict safety protocols for insect handling were applied to further minimize the possibility of any accidental release of transgenic flies. All work on genetically modified organisms was performed under protocols approved by the Institutional Biosafety Committee at Cornell University.

### Genotyping

To obtain genotype information from flies, DNA was extracted by first freezing and then homogenizing individual flies with a 200 μL pipette tip containing 30 μL of solution with 10mM Tris-HCl pH 8, 1mM EDTA, 25 mM NaCl, and 200 μg/mL recombinant proteinase K (Thermo Scientific). The mixture was incubated at 37 °C for 30 min and then 95 °C for 5 min. The solution was centrifuged at 1000g for two min, and 1 μL of the supernatant was used for a PCR reaction, which was purified by gel extraction and assessed via Sanger sequencing. ApE was used to analyze DNA sequence information (http://biologylabs.utah.edu/jorgensen/wayned/ape).

## RESULTS

### Construct design and generation of transgenic lines

We designed two CRISPR/Cas9-based gene drive constructs targeting the X-linked *yellow* gene in *D. melanogaster*. Disruption of this gene causes a recessive yellow (y) phenotype, specified by a lack of dark pigment in adult flies. Our first drive construct contains a Cas9 endonuclease driven by the *nanos* promoter, with a gRNA targeting the coding sequence of the *yellow* gene (Figure 1A). In this case, we expect that most resistance alleles caused by a mutated target site should disrupt the gene (r2 resistance alleles), whereas resistance alleles that preserve the function of *yellow* (r1 resistance alleles) should occur less frequently.

**Figure 1.**
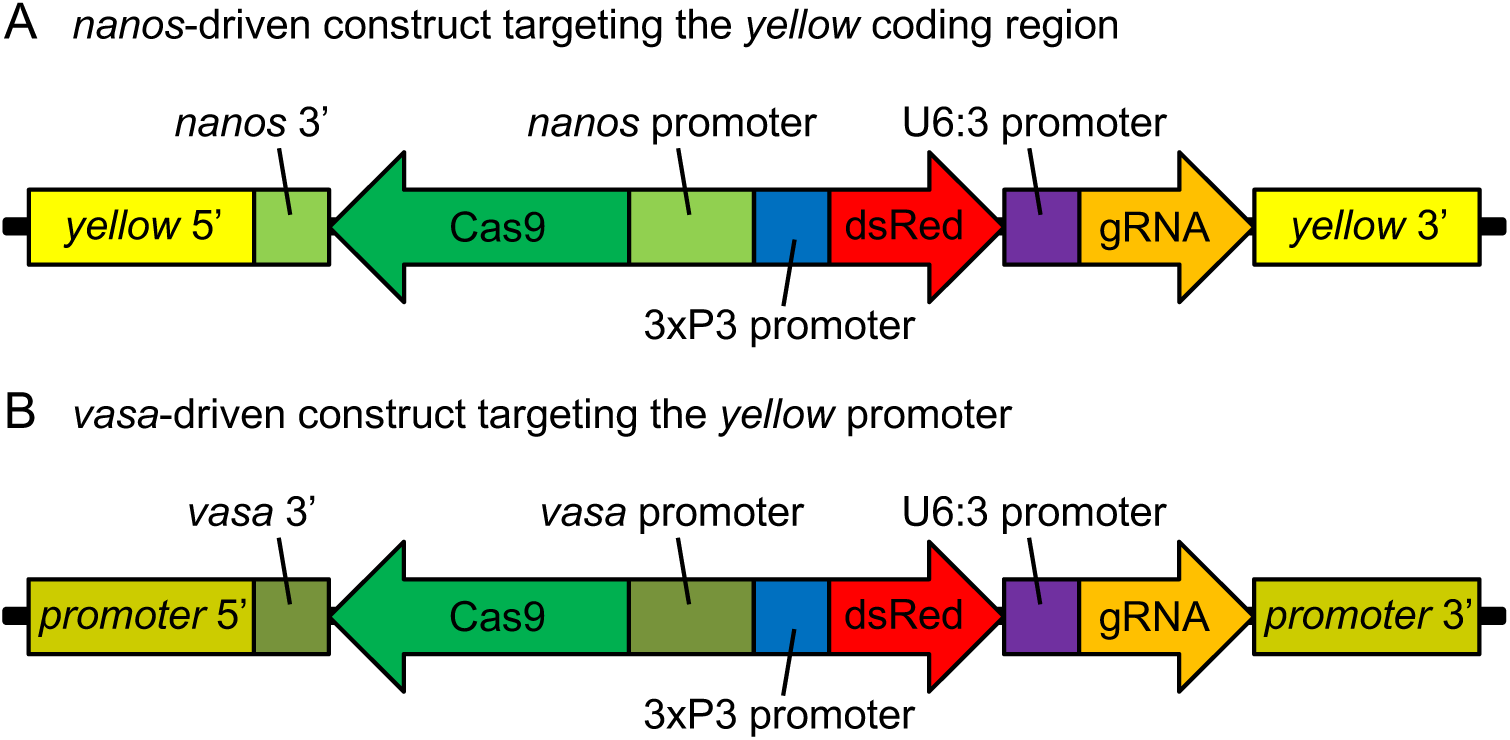
Gene drive constructs. **(A)** Our *namos*-based gene drive construct consists of a Cas9 gene driven by the *namos* promoter and followed by a *namos* 3' UTR. It also contains a dsRed fluorescent marker driven by the 3×P3 promoter, and a gRNA gene driven by the U6:3 promoter. Regions corresponding to the *yellow* gene flank the gene drive components on both ends. **(B)** Our *vasa*/-based gene drive construct has a similar design, but with *vasa* elements replacing the *nemos* ones and flanked by *yellow* promoter sequences.

Our second drive construct contains a gRNA targeting the promoter of the *yellow* gene and a copy of Cas9 driven by the *vasa* promoter (Figure 1B), similar to the construct used in Gantz and Bier^25^. In this case, resistance alleles should be primarily of type r1. The specific insertion site of the *yellow* promoter was selected to induce a (less intense) yellow phenotype in the wings and body when disrupted by a large construct, but to retain male mating success, which is diminished when the *yellow* coding site or downstream regions of the promoter is disrupted by a large construct^35^.

Both of our constructs also encode a dsRed protein, driven by a 3xP3 promoter, which produces an easily identifiable fluorescent eye phenotype (R) that is dominant and allows us to detect the presence of drive alleles (D) in individuals. Box 1 lists the different phenotypes, together with the corresponding genotype combinations in male and female individuals.

**Box 1.**
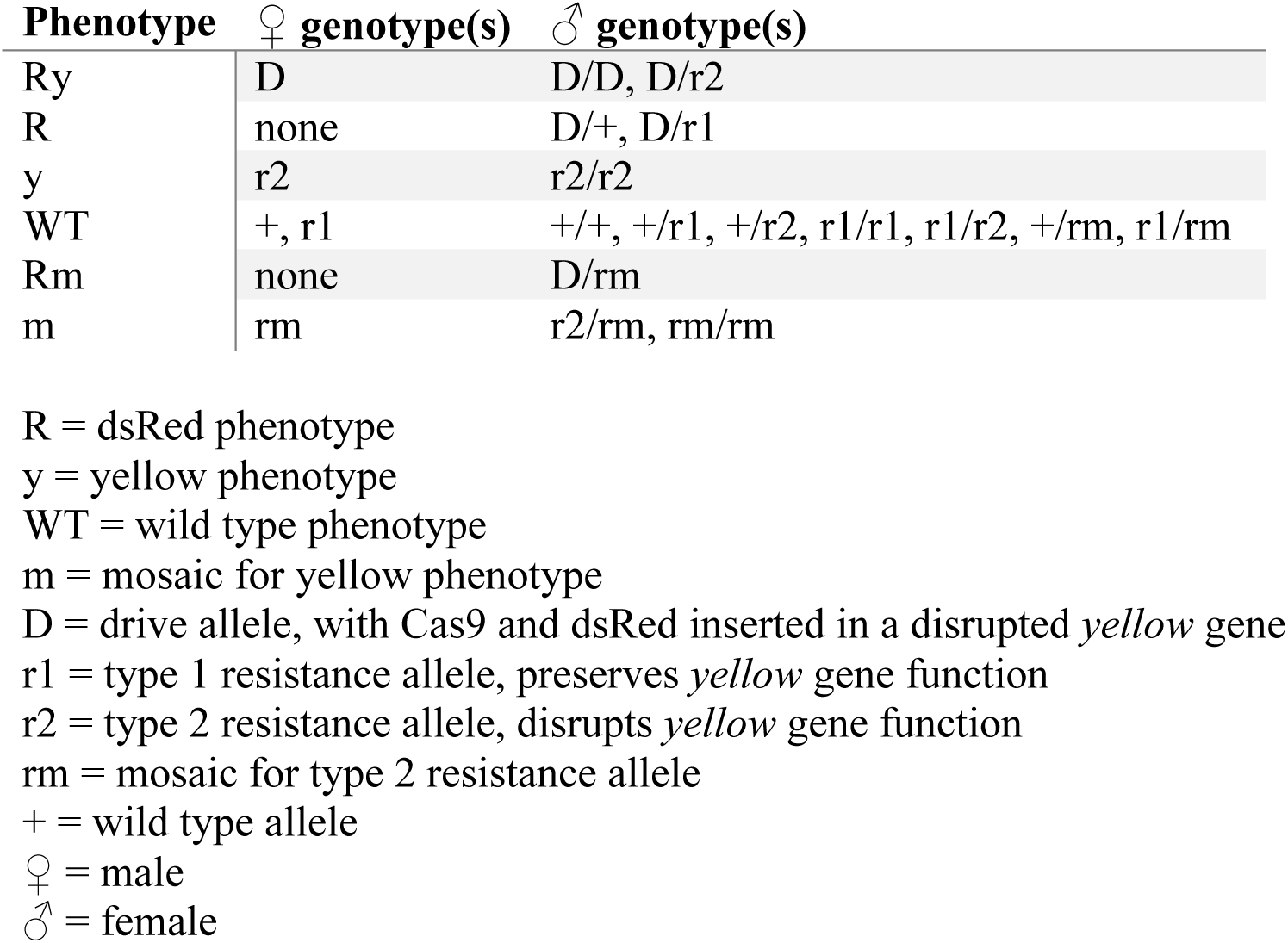
Genotypes and phenotypes.

We generated isogenic fly lines by injecting plasmids containing the constructs into wild type flies from a *w^1118^* line. In both resulting transformed lines, no fluorescent phenotype was immediately visible in injected flies (though some individuals showed yellow mosaicism), but several dsRed transformants were obtained upon mating injected flies to *w^1118^* flies. Although the 3xP3 promoter driving dsRed should be primarily expressed in the eyes and ocelli, we observed expression in the abdomen as well. This enabled the dsRed phenotype to be readily identified in flies with wild type eyes, the pigment of which prevents visualization of dsRed.

### Assessment of drive conversion

To quantify the activity of our gene drive constructs, we performed several crosses between flies from our transgenic lines and the wild type *w^1118^* line. We first verified that when males with the drive allele (genotype D) were crossed with *w^1118^* females (genotype +/+), the progeny followed Mendelian inheritance for an X-linked gene (Tables 1A and 2A). For both constructs, no male progeny and all female progeny from these crosses exhibited the dsRed phenotype, whereas the yellow phenotype was not observed in any progeny, consistent with all daughters having genotype D/+ and all sons having genotype +.

To estimate the drive conversion efficiency of our constructs, we backcrossed the D/+ daughters from the above crosses with males from the *w^1118^* line (genotype +). Under perfect drive conversion (100%), all progeny should then exhibit dsRed phenotype. For the *nanos* drive, we observed dsRed in 81% of the progeny, indicating that only 62.5% of wild type alleles from the D/+ mothers had been successfully converted to drive alleles (Table 1B). For the *vasa* drive, we observed dsRed in 76% of the progeny (Table 2B), corresponding to a conversion rate of 52%, which was slightly lower than that of the *nanos* drive (*p*<0.0001, Fisher’s Exact Test). One *vasa* fly (#3) appeared to have a lower conversion rate (Table S2B, *p*=0.0002, Fisher’s Exact Test), possibly due to leaky vasa-Cas9 expression resulting in a germline mosaic for r1 alleles, so the actual germline conversion rate may be somewhat higher.

**Table 1.**
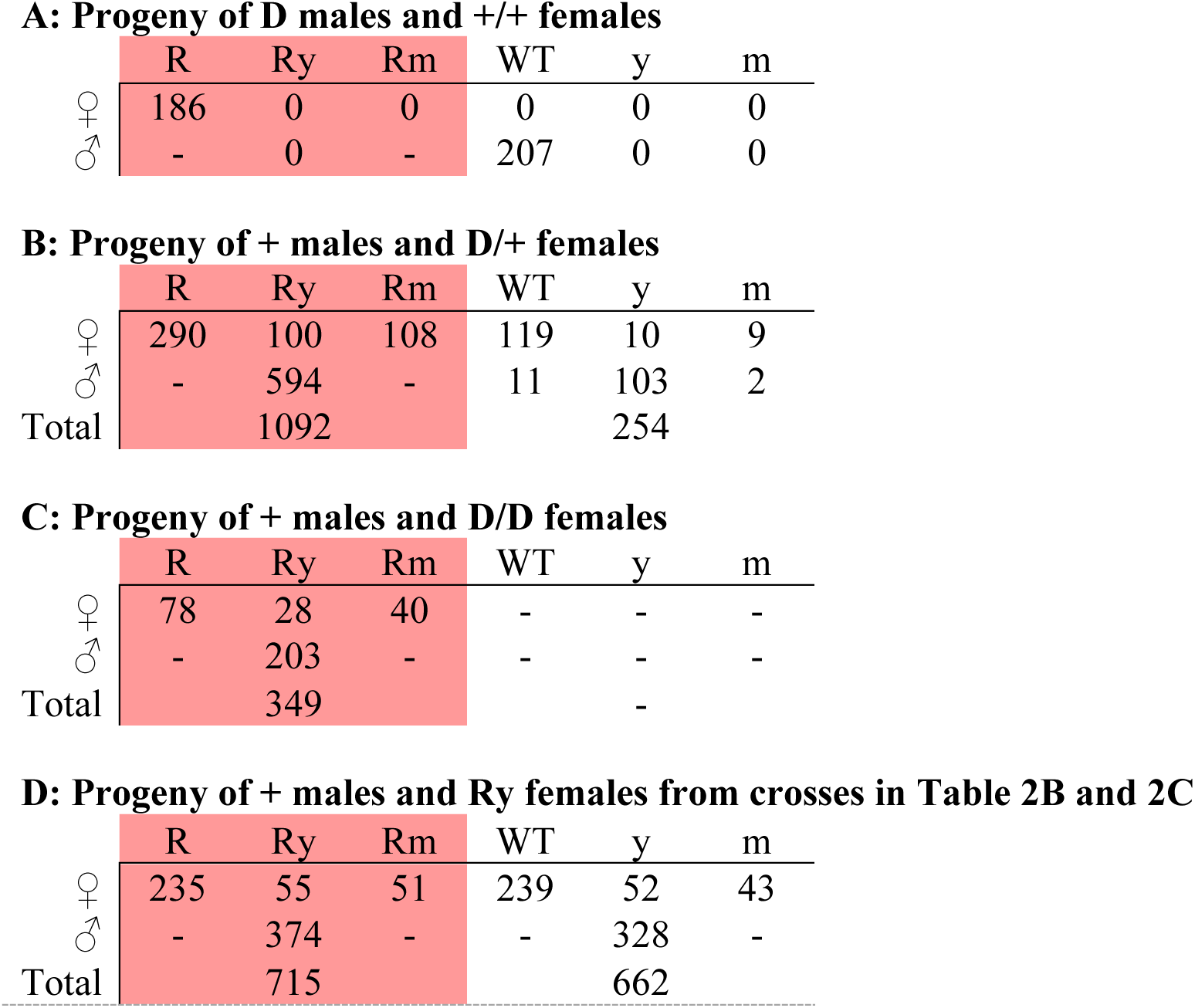
Assessment of *nanos* drive.

**Table 2.**
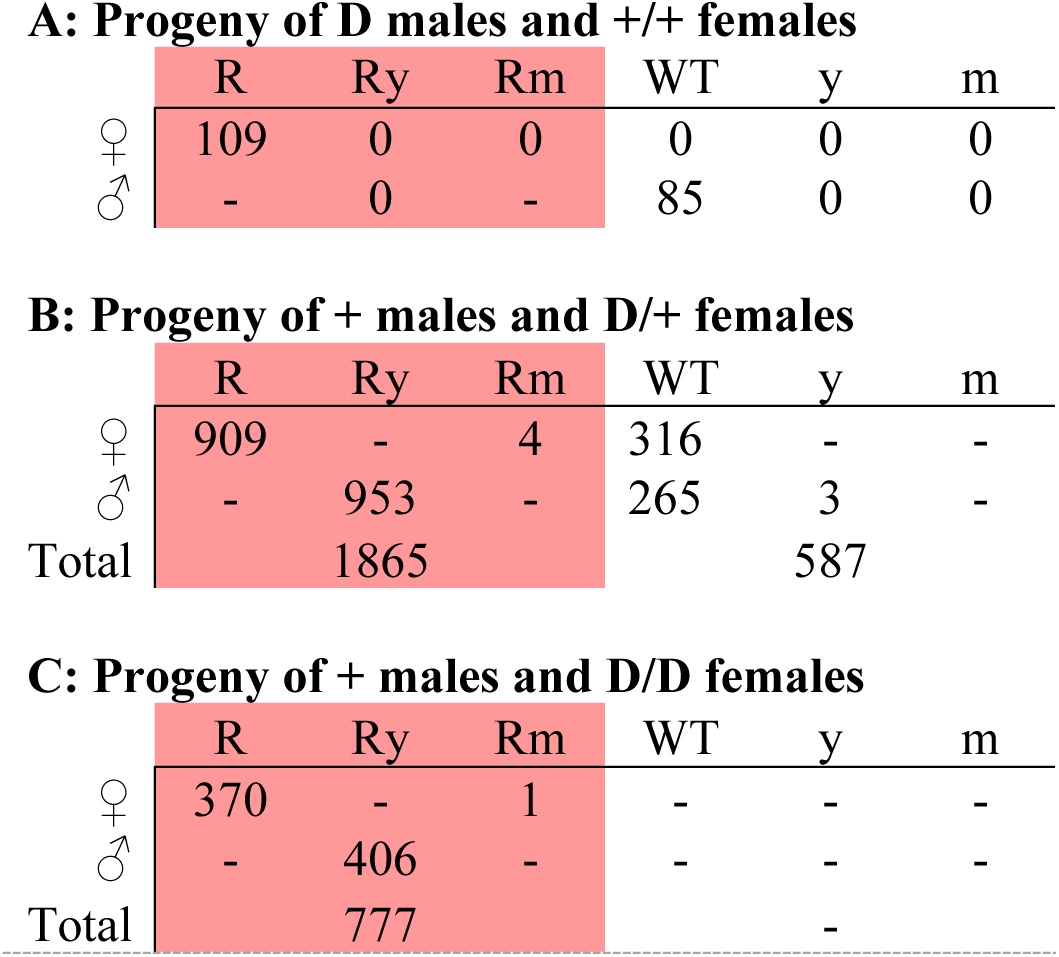
Assessment of *vasa* drive.

When crossing + males and D/D females with the *vasa* drive, all except one daughter were R and all sons Ry (Table 2C), indicating that no significant conversion of wild type alleles to drive alleles took place in the embryo (which would have resulted in daughters with the same Ry phenotype as their D/D mothers). Taken together, these results suggest a different conversion mechanism from a previous study in *D. melanogaster*, which used a similar *vasa* drive targeting *yellow,* but where homing was thought to occur primarily post-fertilization^25^.

### Mechanisms and rates of resistance allele formation

Wild type alleles could, in principle, be converted to resistance alleles via several different mechanisms (Figure 2). For example, resistance alleles could form by NHEJ in the maternal germline, either when Cas9 is expressed before the window for HDR, or a later stage as an alternative to HDR (via partial HDR or NHEJ). They could also form due to persistent Cas9 during meiosis or in a gamete when no template for HDR is available. Alternatively, they could form post-fertilization in the early embryo (though this should occur primarily in daughters for an X-linked homing drive). The results from our crosses, combined with sequencing of resistance alleles, allow us to distinguish some of these scenarios and estimate their relative contributions to the overall rate of resistance allele formation.

**Figure 2.**
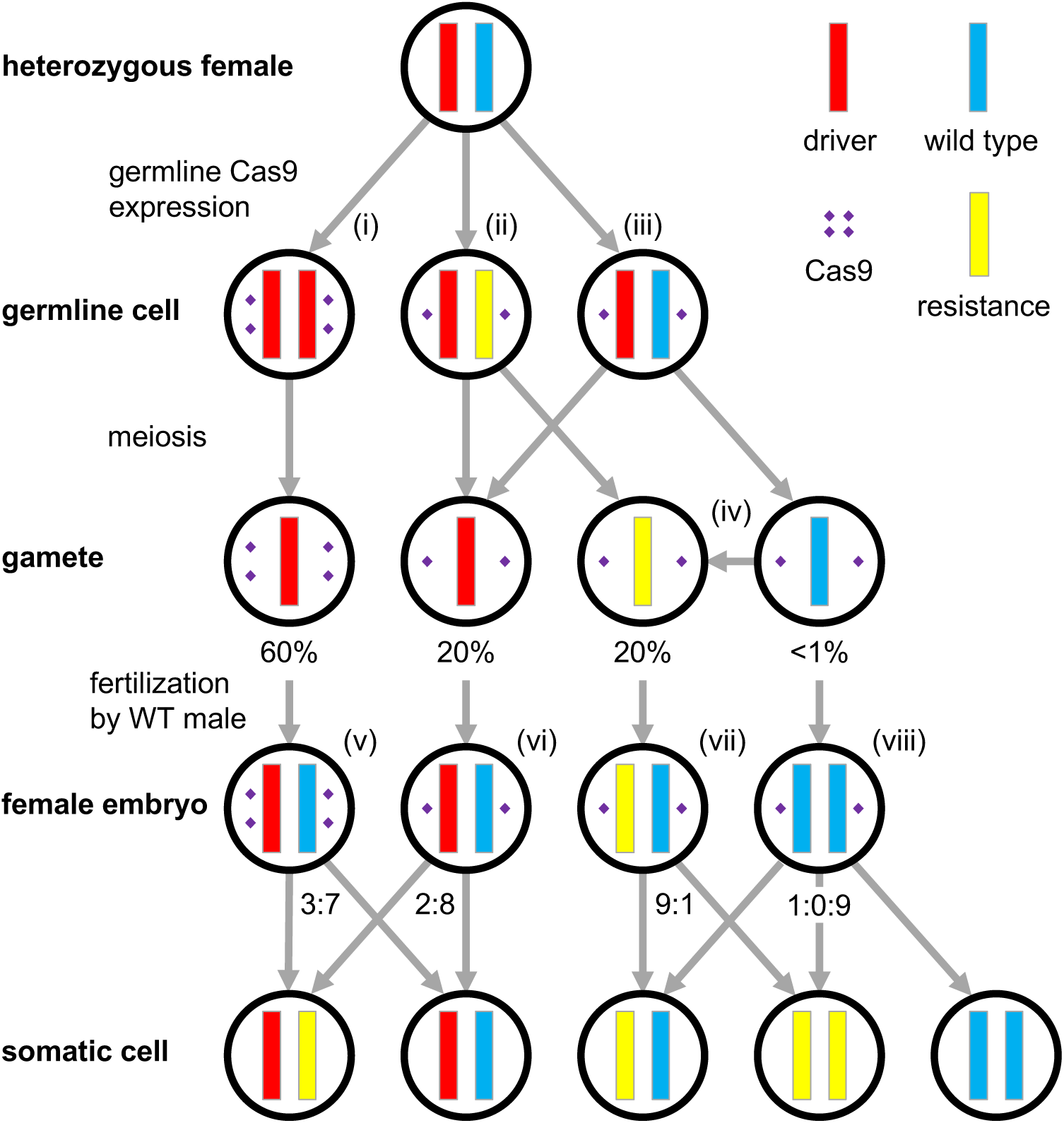
Mechanisms and rates of resistance allele formation. In a heterozygous female with genotype D/+, expression of Cas9 in a germline cell can produce one of three outcomes: (i) successful conversion of the wild type allele into a drive allele by HDR, (ii) formation of a resistance allele when HDR is incomplete or cleavage is repaired by NHEJ, or (iii) continuing presence of the wild type allele if no cleavage occurred or was perfectly repaired. For our *nemos* construct in the *w^1118^* line, we observed successful germline conversion in D/+ females at a rate of approximately 60%, leaving 80% of gametes with gene drive alleles. Almost all remaining gametes (20%) contained resistance alleles, with only less than 1% of gametes carrying a wild type allele, (iv) Note that some of these resistance alleles could have formed during later stages of meiosis or in the gamete when persistent Cas9 cleaved the target while there was no template available for HDR. (v) We also observed the formation of resistance alleles in early female embryos after fertilization by a wild type male, suggesting post-fertilization activity of maternal Cas9 after which cleavage was repaired by NHEJ despite the presence of a template available for HDR. In those embryos that originated from a D/+ or D/D mother, we observed such postfertilization formation of a resistance allele one the paternal chromosome in approximately 30% of embryos, (vi) In embryos that originated from D/r2 mothers, we observed this in only 20% of cases, consistent with presumably lower Cas9 levels in the eggs that would also be found in the r2 gametes from D/+ mothers in the figure. (vii) Formation of resistance alleles was also observed in embryos that did not receive any copy of a drive allele, although formation rates may be lower in this case. (viii) A small number of embryos that inherited wild type alleles from both parents may even have experience double cleavage to form two resistance alleles. Note that any formation of resistance alleles in the embryo may result in mosaicism of adult individuals, as we frequently observed in our crosses. Tables S1–S3 provide the calculations in which these rates were inferred from the phenotypes of progeny from our crosses.

For the *nanos* construct, 103 out of 697 (14.5%) male progeny from the above cross of D/+ females with + males exhibited yellow but not dsRed phenotype (Table 2B), indicative of the presence of r2 resistance alleles. This corresponds to a rate of 29% at which the wild type alleles from D/+ mothers were converted into r2 alleles.

Sequencing of wild type and yellow male progeny from this cross revealed that most of these resistance alleles contained small indels at the gRNA target site, but we also observed two instances of partial HDR, one continuing exactly up to the target sequence of the gRNA gene, which may have enabled repair based on small homologies at each end of the double-strand break (Figure S1A). Of the thirteen wild type males sequenced, ten possessed r1 alleles, and the remaining three had wild type alleles (Figure S1A). These r1 alleles all preserved the reading frame of the *yellow* gene, though there were also several cases of r2 alleles that preserved the reading frame. Based on the phenotype of male offspring, we observed that 97% of wild type alleles in D/+ females were converted to either drive or r2 alleles. Since approximately 10 out of 13 of the remaining alleles contained an r1 resistance allele, this means that less than 1% of alleles passed on to offspring remained wild type, indicating that the *nanos* construct has a high germline cleavage rate in our system.

For the *vasa* drive, out of 29 sequenced sons with wild type phenotype from the cross of D/+ females with + males, all were found to possess r1 alleles (Figure S1B). One additional yellow phenotype male with an r2 allele from the same cross was also sequenced and found to contain a small deletion (interestingly this deletion was shorter than those observed in several r1 alleles). Since *vasa* drive conversion efficiency was 52%, this implies that 48% of wild type chromosomes were converted into resistance alleles.

Several resistance alleles were found to have identical sequences, even for resistance alleles with complicated indels (Figure S1). While the alleles originating from the *vasa* constructs appeared randomly distributed, four out of five of the identical alleles from the *nanos* construct were found in flies that shared the same batch of the parents. These data support the idea that resistance alleles in the *nanos* drive could have formed in early germline stem cells that eventually gave rise to multiple progeny, though it does not rule out the formation of resistance alleles by other pathways as well.

We also found strong evidence for post-fertilization formation of resistance alleles in a fraction of female embryos. In our crosses of *nanos* drive D/+ females with + males, 20% of the daughters that received the drive were Ry (Table 1B), suggesting that the *yellow* gene copy inherited from the wild type father must have been disrupted as well. A similar number of progeny (22%) were mosaic for yellow phenotype, indicating that Cas9 cleavage and NHEJ repair was delayed until after the embryo had undergone several divisions, thus resulting in only some cells experiencing cleavage and r2 allele formation. The proportion of daughters with yellow phenotype was higher among those individuals that also exhibited dsRed than those who did not (*p*=0.0002, Fisher’s Exact Test). This suggests that successfully converted drive alleles either expressed additional Cas9 prior to meiosis, which persisted into the embryo, or that Cas9 was actively expressed after meiosis.

When + males were crossed with nanos-drive D/D females, all offspring showed dsRed phenotype, yet 19% of daughters again also exhibited the yellow phenotype with an additional 27% mosaic (Table 1C). The fraction of these Ry females among all females with dsRed phenotype was similar to that observed in the cross between + males and D/+ females. This implies that most maternal Cas9 persisting to the embryo stage was expressed after drive conversion events.

Daughters with Ry phenotype from the previous two crosses could either be genotype D/D or D/r2, while daughters with R phenotype could either be D/+ or D/r1. Such r1 resistance alleles are indistinguishable from wild type alleles by phenotype, yet we could still detect their presence via their inability to be successfully converted into drive alleles when several R phenotype daughters were mated to + males and their progeny scored. In cases of normal levels of conversion, progeny were included in Table 1B. However, one fly showed no germline conversion and thus, likely possessed an r1 allele in addition to its drive allele.

To detect the presence of r2 alleles, we crossed the Ry daughters with + males. We observed dsRed in 52% of the resulting progeny (Table 1D), suggesting that most of the Ry mothers were D/r2, since we would expect 100% to have a dsRed phenotype if mothers had been D/D. The progeny of this cross also contained 16% yellow daughters, with an additional 14% mosaic. The numbers of these were similar regardless of whether a drive allele was inherited, implying that Cas9 persisted through to the embryo after maternal expression in early oocytes rather than being expressed after meiosis.

The rate of total embryonic cleavage events that formed r2 alleles in the progeny of D/r2 females (including both full yellow and mosaic females) was significantly lower than in the progeny of the combined D/D and D/+ females (Fisher’s Exact Test, *p*=0.0001), possibly because embryos from the D/D mothers had higher Cas9 levels due to their second D allele copy. Sequencing of ten of these D/r2 females revealed that all were genetically mosaic for 2-4 different r2 resistance allele sequences, indicating that Cas9 cleavage occurred at a high rate early in the embryo, but usually not in the zygote.

Post-fertilization formation of resistance alleles in the embryo was also observed for the *vasa* drive, as evidenced by the finding that a small number of daughters from the cross between D/+ females and + males had yellow or mosaic phenotype (Table 2B). A fraction of resistance alleles must therefore have disrupted the *yellow* promoter sufficiently to reduce expression of *yellow* in the wings and body.

When crossing several of these R daughters with + males, most progeny showed normal levels of drive conversion and were included in Table 2B. However, r1 alleles were apparent when only 50% of offspring contained a drive allele (Table S2B) instead of the 76% expected from D/+ fly. Some flies appeared to have drive inheritance significantly different from both 50% and 76% (*p*<0.01, Fisher’s Exact Test), which may indicate mosaicism within the germline due to leaky vasa-Cas9 expression.

### Differences in drive efficiency and resistance rates between distinct genetic backgrounds

Drive efficiency and rates of resistance allele formation could in principle depend on the specific genetic background of an individual, for example due to genetic effects on Cas9 expression levels or the efficiency and fidelity of different repair pathways. To test for such potential differences, we studied our *nanos* construct in several genetically distinct fly lines, including the Canton-S line and five lines from the Global Diversity Lines^32^, one from each continent. We also studied the *vasa* construct in the Canton-S line.

Males with the drive construct were first crossed to females from the Canton-S and Global Diversity Lines. Offspring were + males and D/+ females. These were allowed to mate, and their progeny were phenotyped. Overall, we observed somewhat lower conversion efficiency for the *nanos* construct in each of these lines than for our original *w^1118^* line. Specifically, we found that conversion efficiency in D/+ mothers varied between 40-55% in the 6 lines (Table 3A). We also found significant differences for rates at which r2 alleles were formed, which ranged between 35% and 52% in the germline of D/+ females (detected in y phenotype sons).

The most marked differences between lines were found in the rate of post-fertilization formation of r2 alleles in the early embryo of daughters, which ranged between 4% and 56%. The *vasa* construct in the Canton-S line also had significantly lower drive conversion efficiency than the *w^1118^* line (Table 3B). These differences between lines demonstrate that genetic differences between individuals can indeed have a substantial impact on drive efficiency and resistance allele formation rates.

**Table 3.**
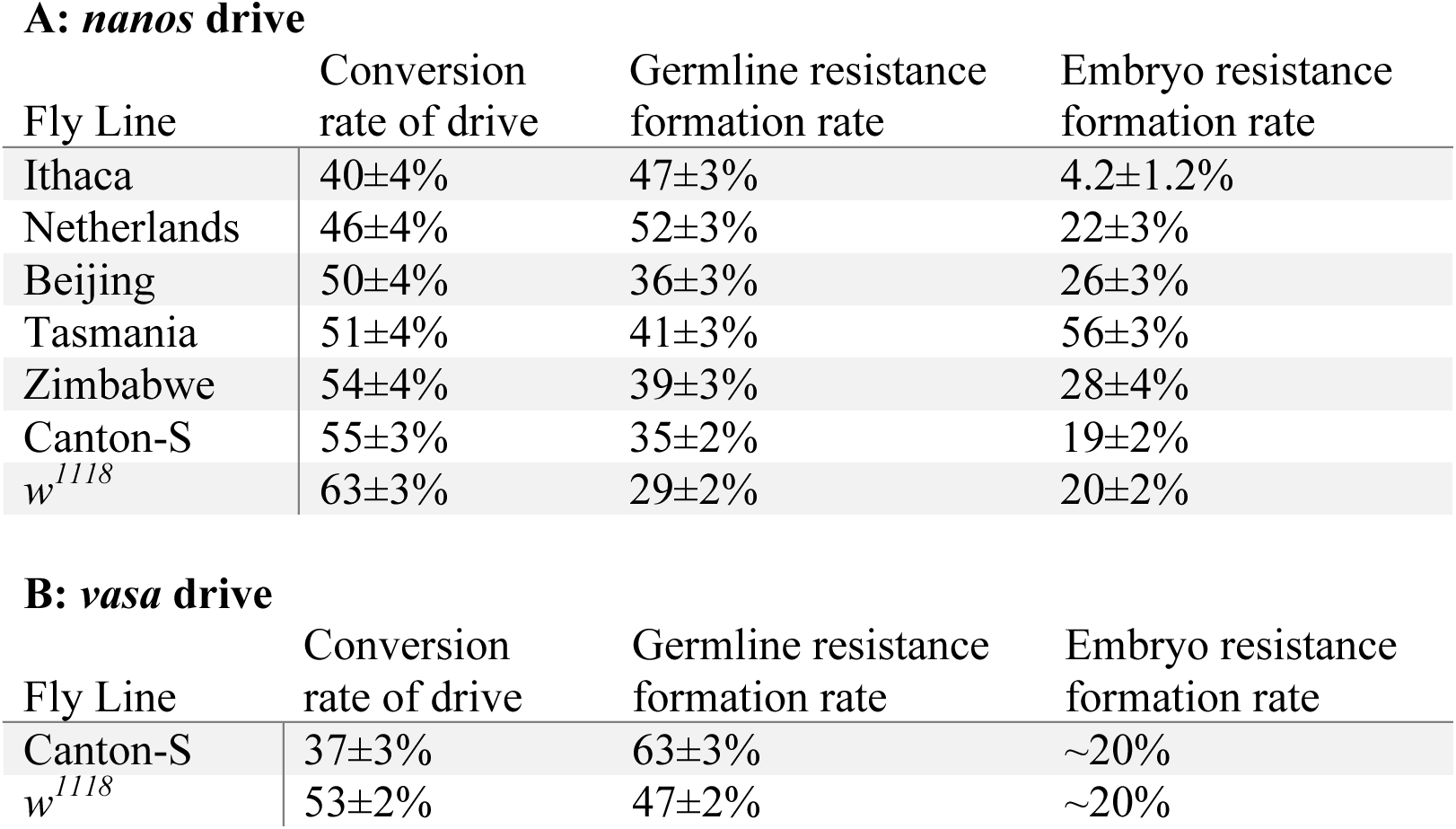
Drive parameters for several *D. melanogaster* lines.

These data do not yet allow us to infer whether genetic variation impacting efficiency is distinct from that impacting the formation of post-fertilization resistance alleles, although the two do not appear to be correlated. On the other hand, drive efficiency appears to be inversely related to the formation of resistance alleles in the maternal chromosome (Figure S3, R^2^=0.89), lending support to the idea that 97-100% of all maternal wild type alleles from D/+ heterozygotes were typically converted to either a drive allele or a resistance allele.

## DISCUSSION

The idea of using CRISPR gene drives for genetic modification of entire species has ignited an intense debate about potential applications as well as risks of such approaches^36–38^. While some consider gene drives a means for solving important global challenges, such as the fight against vector-borne diseases or the preservation of endangered species, others are alarmed by the possibility of unintended consequences. In this study, we highlight the fact that to make CRISPR gene drives at least technically feasible, key obstacles still need to be overcome.

We focused specifically on the issue of resistance against a drive, which should evolve when cleavage-repair by NHEJ produces mutated target sites that are no longer recognized by the drive’s gRNA. To study this process, we designed two drive constructs in *D. melanogaster* (using the *nanos* and *vasa* promoters) following a similar design as a previous study^25^, with modifications that allowed for improved assessment of drive efficiency and resistance allele formation. Both of our constructs produced resistance allele at high rates, consistent with previous experiments ^25,27,28^. The specific design of our constructs allowed us to determine where in the life cycle these resistance alleles arose, revealing formation both prior to fertilization in the germline as well as post-fertilization in the embryo due to maternally deposited Cas9 (Figure 2). Additionally, we found that resistance allele formation rates vary markedly depending on the specific genetic background of the individual flies.

Homing and integration of our constructs appeared to occur in the germline. This differs from a previous study in *D. melanogaster,* which used the same *yellow* target site as our *nanos* construct and a *vasa* promoter identical to the one in our own *vasa* construct^25^, yet where homing was thought to occur in the early embryo^25^. However, an alternative interpretation is that resistance alleles that also disrupted *yellow* formed post-fertilization in their experiments, which gave the appearance of much higher drive efficiency. This system also produced yellow daughters of males with the homing drive, which may be due to high levels of leaky expression of Cas9 from the *vasa* promoter. We saw such leaky expression to a more limited extent in our *vasa-based* system (Tables S2B and S2D), likely due to the localization of our *vasa* drive to the *yellow* promoter instead of the coding sequence.

A later study involving the same research group used a homing drive in *Anopheles stephensi* mosquitoes with a very similar mechanism to our *D. melanogaster* drives^28^. In this drive, which utilized the *A. stephensi vasa* promoter to express Cas9, drive conversion efficiency was higher than for either of our constructs, with a correspondingly lower rate of resistance alleles formed in the germline. However, the rate of resistance alleles formed post-fertilization in the embryo was also much higher. These characteristics could potentially be caused by differences in the level of Cas9 expression, with higher expression in the mosquito system resulting in greater initial homing efficiency, but also greater Cas9 persistence in embryos causing correspondingly higher post-fertilization resistance allele formation. The differences in resistance allele formation may lend support to models in which resistance alleles form prior to or after the window for HDR, rather than as an alternative to successful drive conversion.

One of our constructs utilized the *nanos* promoter in an attempt to reduce the formation of resistance alleles due to its germline-restricted expression compared to the *vasa* promoter, which has leaky somatic expression. While resistance alleles still formed post-fertilization for our *nanos* drive, this occurred at the same rate for insects that inherited the drive allele as those that did not (Table 2C). This contrasts with the *A. stephensi* system using the *vasa* promoter^28^, in which more resistance alleles were formed post-fertilization when the drive was inherited compared to when a resistance allele was inherited. We also found some instances of leaky postfertilization expression with this promoter (Table S2C). Thus, the *nanos* promoter appears to offer a modest advantage in terms of reducing formation of resistance alleles, as well as potentially slightly higher drive efficiency in our system, though the latter may be because our *nanos* system contained a Cas9 that was codon-optimized for insects compared to our *vasa* system that contained a Cas9 codon-optimized for mammalian expression. It could also be an artifact of small levels of leaky vasa-induced formation of resistance alleles in the germline of flies thought to be entirely D/+.

Theoretical studies have shown that to effectively spread in a population, a gene drive system requires significantly lower rates of resistance allele formation than observed among current Cas9-based drives, including those developed in this study^29,30^. This is particularly relevant for approaches aimed at population suppression, which will probably be thwarted by any measurable rate of resistance allele formation. However, population modification approaches are also sensitive to the evolution of resistance if the drive allele carries even a small fitness cost, which is likely due to the presence of a payload gene, off-target cleavage effects of Cas9, or even the expression of the large Cas9 protein itself.

As we have highlighted in this study, one prerequisite for lowering resistance potential is better control of Cas9 expression. An ideal promoter would offer the high rate of drive conversion seen for the *vasa* promoter in the *A. stephensi* system^28^, but with lower persistence of Cas9 to the embryo and no leaky expression. While resistance allele formation in the embryo could potentially be lowered by using an autosomal drive that functions only in males, resistance alleles that form prior to fertilization in the germline would remain problematic.

Several additional strategies have been suggested to lower resistance potential. Those involving the use of multiple gRNAs targeting different sites have drawn a considerable amount of attention^3,4,39^. Yet, with the current rates of resistance allele formation, they are unlikely to be effective alone and would probably need to be combined with other approaches such as a better promoter. Resistance allele formation could also be suppressed by utilizing a shRNA gene as part of the drive designed to suppress NHEJ in the germline and early embryo^40^. Another possibility is to target a haplolethal gene such that its function is preserved only under successful conversion^3,4,39^. However, rare resistance alleles that also preserve gene function could still prove difficult to overcome for such an approach. A haplosufficient gene could be targeted in a similar manner, which may enable the gene drive to persist longer in a population due to lower fitness costs of the drive compared to resistance alleles that disrupt the target gene function. Either of these strategies might be considerably enhanced with the use of multiple gRNAs because of increased potential to disrupt the gene of interest, even if HDR does not take place.

Whether a gene drive can successfully spread in the wild ultimately depends on how it performs in a genetically diverse population. To study this question, we tested our *nanos* drive in five different genetic backgrounds drawn from the *D. melanogaster* Global Diversity Lines and also the wild type Canton-S laboratory strain. While conversion efficiency and germline resistance allele formation showed only modest variation across lines, post fertilization resistance formation in the embryo was significantly heterogeneous, varying over more than an order of magnitude. This suggests that resistance rates are not a fixed feature of a given gene drive construct, but can vary substantially between individuals and genetic backgrounds. Future studies will attempt to identify the genetic loci responsible for this variation.

The variability in resistance rates we observed in this study has important implications for the feasibility of gene drive strategies in the wild. The likelihood that resistance evolves against a drive should be determined primarily by those individuals that have a high rate of resistance allele formation, even when the average rate in the population is low. This will be particularly relevant for target populations that harbor high levels of genetic diversity, such as *A. gambiae^41^.* It also has important implications for the assessment of gene drive parameters in the laboratory, which should include cage experiments of large, genetically diverse populations followed over many generations, instead of focusing on crosses of isogeneic lines. Finally, variation in drive parameters between individuals will need to be included in our theoretical models, which currently rely on rather simplistic assumptions such as constant resistance and conversion rates across the whole population^6–11,29,30^.

While certain gene drive strategies may be able to tolerate a low level of resistance allele formation, for instance if they only require the drive allele to persist in the population for a limited period of time, the specific outcome of such strategies still depends strongly on the fitness costs of the payload and the drive itself^29,30^. The assessment of such fitness costs therefore provides another important avenue of future research. Extensive modeling efforts will be required to determine what levels of these parameters may be acceptable to retain efficiency for different types of drives in different scenarios.

## ACKNOWLEDGEMENTS

We are grateful to Kate O'Connor-Giles, Melissa Harrison, Jill Wildonger, and Simon Bullock for providing plasmids. This research was supported by startup funds from the College of Agriculture and Life Sciences at Cornell University to P.W.M.

## SUPPLEMENTARY INFORMATION

### Supplementary methods

The following tables show the DNA fragments used for Gibson Assembly of the listed plasmids. PCR products are shown with the oligonucleotide primer pair used, and plasmid digests are shown with the restriction enzymes used.

|  |  |  |  |
| --- | --- | --- | --- |
| **IHDyi1** | *Template* | *Oligo/Enzyme 1* | *Oligo/Enzyme 2* |
| *PCR Product* | Genomic DNA | YellowLeft_F | YellowLeft_R |
| *Plasmid Digest* | pDsRed-attp | EcoRI-HF | NruI-HF |
| **IHDypi1** | *Template* | *Oligo/Enzyme 1* | *Oligo/Enzyme 2* |
| *PCR Product* | Genomic DNA | YellowPLeft_F | YellowPLeft_R |
| *Plasmid Digest* | pDsRed-attp | EcoRI-HF | NruI-HF |
| **IHDyg1** | *Template* | *Oligo/Enzyme 1* | *Oligo/Enzyme 2* |
| *PCR Product* | none | Yellow_gRNA_F | Yellow_gRNA_R |
| *Plasmid Digest* | pCFD3 | BbsI | none |
| **IHDypg1** | *Template* | *Oligo/Enzyme 1* | *Oligo/Enzyme 2* |
| *PCR Product* | none | YellowP_gRNA_F | YellowP_gRNA_R |
| *Plasmid Digest* | pCFD3 | BbsI | none |
| **IHDyi2** | *Template* | *Oligo/Enzyme 1* | *Oligo/Enzyme 2* |
| *PCR Product* | IHDyg1 | YellowU6_3_gRNA1_F | YellowU6_3_gRNA1_R |
| *PCR Product* | Genomic DNA | YellowRight_F | YellowRight_R |
| *Plasmid Digest* | IHDyi1 | PstI-HF | SpeI-HF |
| **IHDypi2** | *Template* | *Oligo/Enzyme 1* | *Oligo/Enzyme 2* |
| *PCR Product* | IHDypg1 | YellowU6_3_gRNA_F | YellowPU6_3_gRNA_R |
| *PCR Product* | Genomic DNA | YellowPRight_F | YellowPRight_R |
| *Plasmid Digest* | IHDypi1 | SpeI-HF | XhoI |
| **IHDyN1** | *Template* | *Oligo/Enzyme 1* | *Oligo/Enzyme 2* |
| *PCR Product* | pnos-Cas9-nos | NosCas9_1_F | NosCas9_1_R |
| *PCR Product* | pnos-Cas9-nos | NosCas9_2_F | NosCas9_2_R |
| *Plasmid Digest* | IHDyi2 | NotI-HF | SphI-HF |
| **IHDypV1** | *Template* | *Oligo/Enzyme 1* | *Oligo/Enzyme 2* |
| *PCR Product* | pvasa-Cas9 | VasaCas9_1_F | VasaCas9_1_R |
| *PCR Product* | pvasa-Cas9 | VasaCas9_2_F | VasaCas9_2_R |
| *Plasmid Digest* | IHDypi2 | NotI-HF | SphI-HF |

### Construction primer list

`YellowLeft_F:AATTAACCAATTCTGAACATTATCGCCTAGGGTTGCGAGGTTTTAGGACTGAAAGAGCAC`

`YellowLeft_R:TGCATATGTCCGCGGCCGCTAGCATGCAAGAATTCTTCCAGTGTCCAAAACCCACAGCC`

`YellowPLeft_F:CCAATTAACCAATTCTGAACATTATCGCCTAGGGCGCTCCGCTTGATGTTGTTTTGTT`

`YellowPLeft_R:TATGTCCGCGGCCGCTAGCATGCAAGAATTCTGAGGGTCAAATATTTGGTTTCCGCTAGT`

`Yellow_gRNA_F:TATATATAGACCTATTTTCAATTTAACGTCGGTTTTGGACACTGGAACCG`

`Yellow_gRNA_R:ATTTTAACTTGCTATTTCTAGCTCTAAAACCGGTTCCAGTGTCCAAAACC`

`YellowP_gRNA_F:TATATATAGACCTATTTTCAATTTAACGTCGATGATTCACAATTCACTGA`

`YellowP_gRNA_R:ATTTTAACTTGCTATTTCTAGCTCTAAAACTCAGTGAATTGTGAATCATC`

`YellowU6_3_gRNA_F:ATGTATGCTATACGAAGTTATAGAAGAGCACTAGTTTTTTTGCTCACCTGTGATTGCTC`

`YellowU6_3_gRNA_R:CGATGCCCACGGACGCGTCCTGCAGGATGCATACGCATTAAGCGAACATT`

`YellowRight_F:GCAT CCTGCAGGACGCGT CCGTGGGCATCGGCAATACCACC`

`YellowRight_R:GATTGACGGAAGAGCCTCGAGCTGCACACACAGTGGACTACATTGCCTGAATTGGCGGGC`

`YellowPU6_3_gRNA_R:TTCACAATTCACACGCGTCCTGCAGGATGCATACGCATTAAGCGAACATT`

`YellowPRight_F:GCAT CCTGCAGGACGCGT GTGAATTGTGAATCATCGGTGACGCC`

`YellowPRight_R:AACTCGATTGACGGAAGAGCCTCGAGCACACAGTGCACAAGGATCCACCCTTTGTCCTGG`

`NosCas9_1_F:ACTACGATCGCAGGTGTGCATATGTCCGCGGCCGCGCTTCGACCGTTTTAACCTCGAAAT`

`NosCas9_1_R:TCCTGTATATCGGCGCCTCTT`

`NosCas9_2_F:AAGAGGCGCCGATATACAGGA`

`NosCas9_2_R:GCTGTGGGTTTTGGACACTGGGAATTCTTGCATGCTCCTTCCTGGCCCTTTTCGAG`

`VasaCas9_1_F:ACTACGATCGCAGGTGTGCATATGTCCGCGGCCGCCTGCAGCTGGTTGTAGGTGCAGTTG`

`VasaCas9_1_R:GTTCCTCGGTGCCGTCCATCTTTTC`

`VasaCas9_2_F:GAAAAGATGGACGGCACCGAGGAAC`

`VasaCas9_2_R:GGGTTTTGGACACTGGGAATTCTT GCATGGCTAGCCAACACGAAGAGCAGCAGTGTGGTG`

### Sequencing primer list

`YellowLeft_S_F:AGAGCCATTAGCACGGCAGTTACCA`

`YellowRight_S_R:TCGAATGGGCGAAAGGGACATACCA`

`YelProLeft_S_F:ATTACCCACTTAGGGCACCCCCAAC`

`YelProRight_S_R:CAGTGTTCATCTTTATCGGCGACTGCAA`

**Figure S1A.**
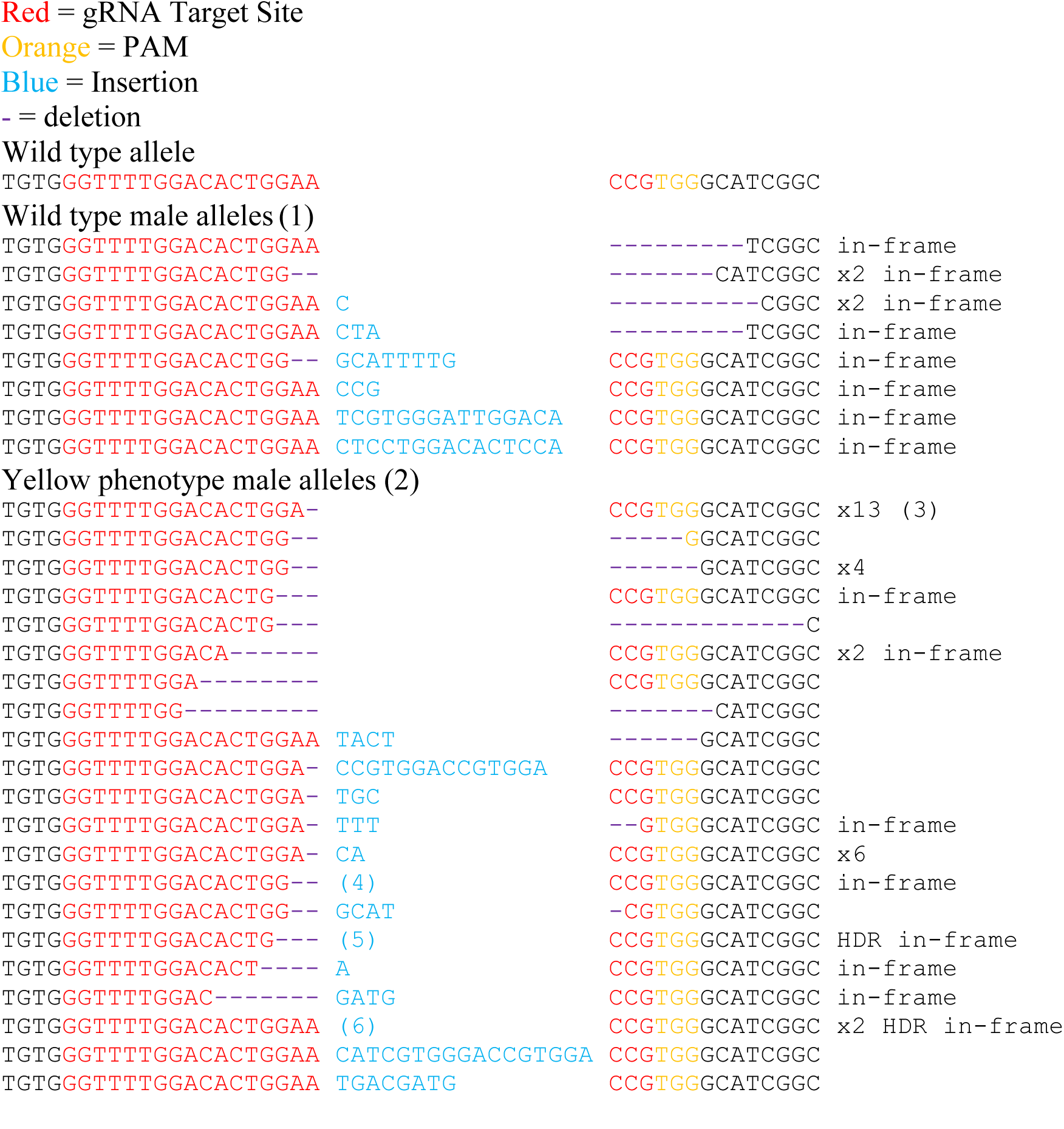
Resistance allele sequences from the *nanos* drive.

**Figure S1B.**
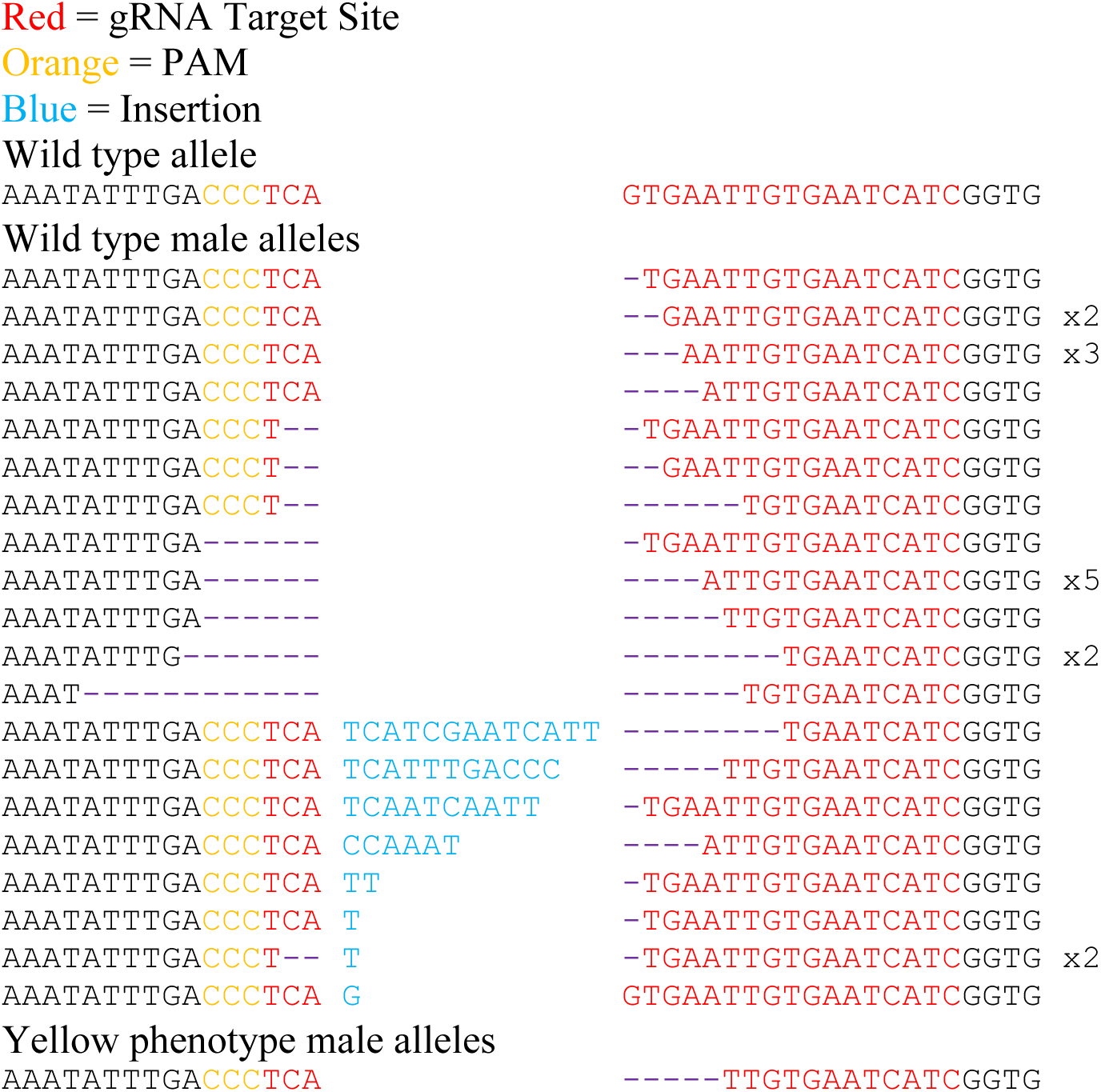
Resistance allele sequences from the *vasa* drive.

**Figure S2.**
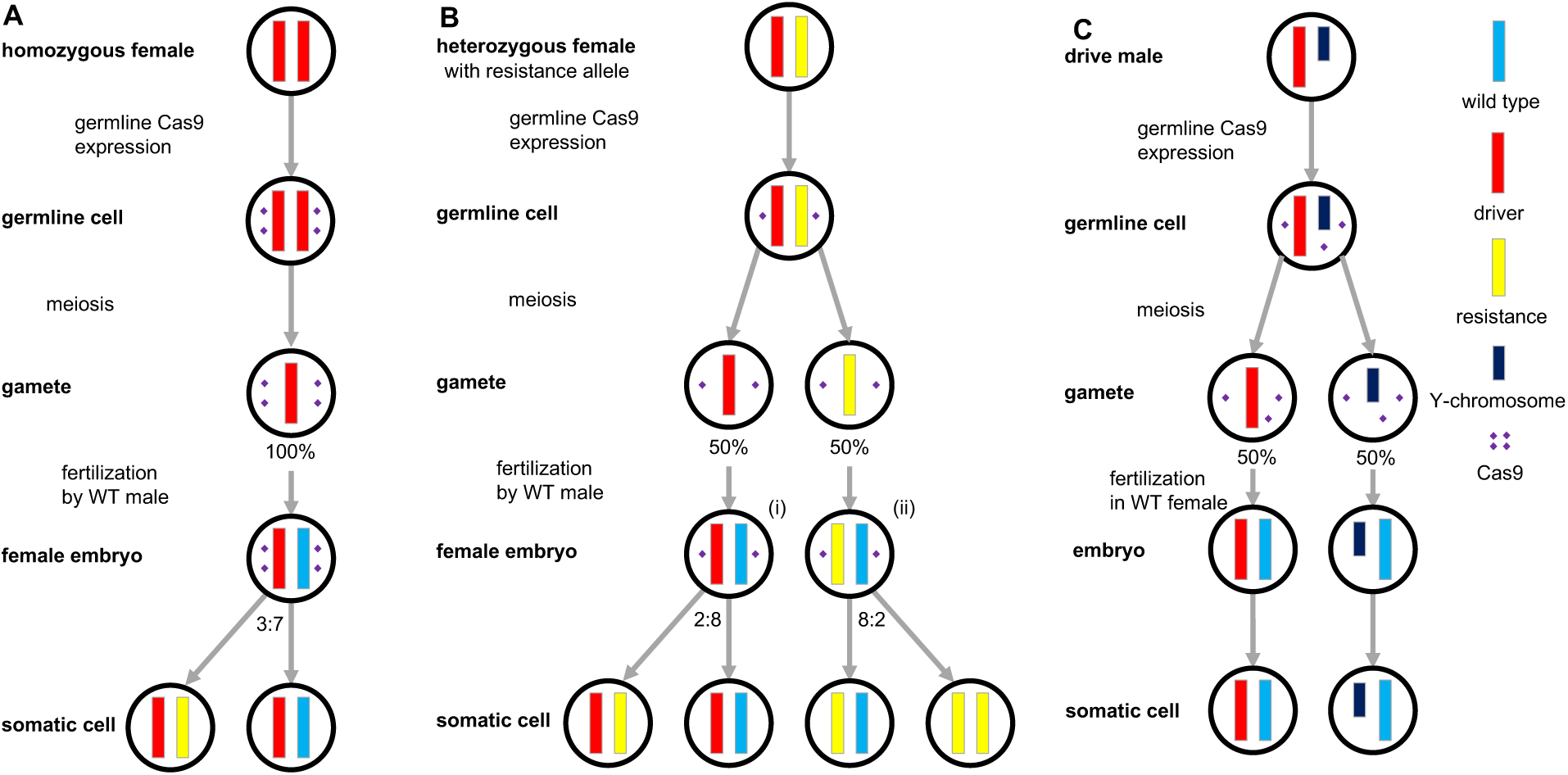
Mechanisms and rates of resistance allele formation in additional genotypes. (**A**) In a homozygous female with genotype D/D, high expression of Cas9 results in relatively high formation of resistance alleles (~30%) in early female embryos after fertilization by a wild type male due to persistence of maternally expressed Cas9. (**B**) In a heterozygous female with genotype D/r2, no drive conversion takes place in the germline, and lower expression of Cas9 results in reduced formation of resistance alleles (~20%) in female embryos after fertilization by a wild type male. (**C**) In a male with the gene drive, no drive conversion takes place in the germline due to the presence of only one X-chromosome. Additionally, the relatively small size of the gamete means that a significant amount of Cas9 does not persist to the embryo, resulting in little to no post-fertilization formation of resistance alleles. Note that any formation of resistance alleles in the embryo may result in mosaicism of adult individuals, as we frequently observed in our crosses. Additionally, leaky expression of Cas9, as observed in the *vasa* construct, may potentially form additional resistance alleles in the embryo or later stages. Tables S1-S3 provide the calculations in which these rates were inferred from the phenotypes of progeny from our crosses.

**Figure S3.**
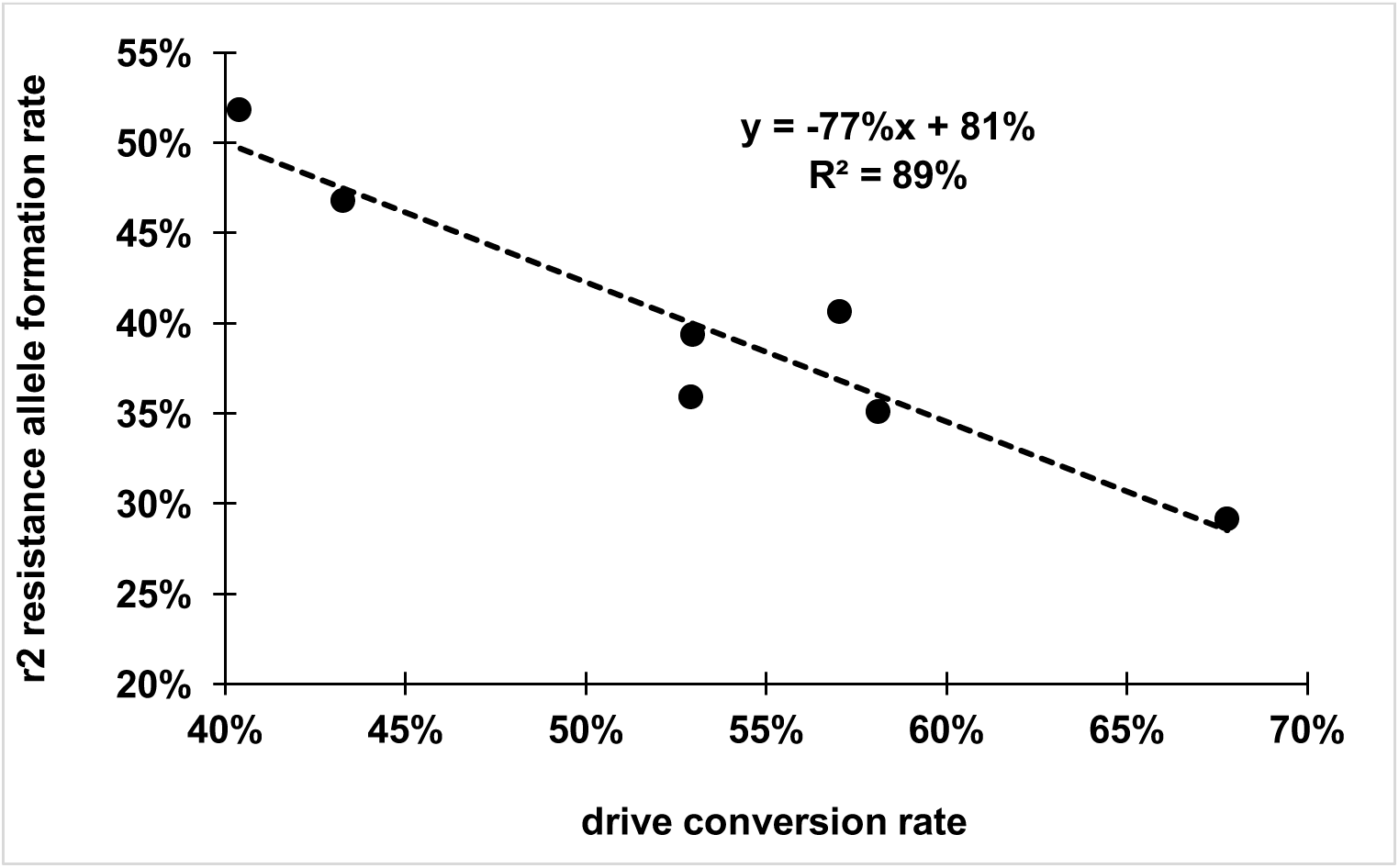
Negative correlation between drive conversion rate and r2 resistance allele formation rate for the *nanos*-drivt across different genetic backgrounds. All rates were obtained from Table S3.

